# Biting time of day in malaria mosquitoes is modulated by nutritional status

**DOI:** 10.1101/2025.04.28.650966

**Authors:** Catherine E. Oke, Samuel S. C. Rund, Maxwell G. Machani, Abdul Rahim Mohammed Sabtiu, Yaw Akuamoah-Boateng, Yaw A. Afrane, Sarah E. Reece

## Abstract

**Background:** Transmission of vector-borne pathogens follows daily rhythms, occurring at the time of day that vectors forage for blood. Control measures such as insecticide-treated bed nets (ITNs) have been particularly successful for reducing malaria transmission, because they exploit the nocturnal biting behaviour of the *Anopheles* spp. that vector malaria. However, shifts in biting behaviour to earlier or later hours when people are unprotected can undermine the efficacy of ITNs. Despite the implications for malaria transmission, the mechanisms underlying these shifts remain poorly understood. Because food availability mediates activity and foraging rhythms, and ITNs block access to human blood as a food source, we hypothesized that nutritional deprivation could cause mosquitoes to shift their biting behaviour to earlier or later in the diel cycle.

**Methods:** We provided female *Anopheles gambiae s.l.* mosquitoes with a blood meal on day 3 post-emergence, and access to one of three feeding treatments that varied in nutritional resources: (i) 0.5% sucrose, (ii) 10% sucrose, or (iii) 10% sucrose plus an additional blood meal on day 6. We released mosquitoes into a semi-field system on day 10 with human-mimic traps to investigate how food availability impacted the time of day that mosquitoes host seek.

**Results:** Nutritional resources determine both the likelihood and time of day that host seeking occurs. Specifically, low-resourced mosquitoes were 2-3 fold more likely to host seek overall, and 5-10 fold more likely to host seek at an earlier time of day than well-resourced mosquitoes (fed 10% sucrose with and without an additional blood meal), which predominantly sought a host in the second half of the night time.

**Conclusions:** We reveal that mosquito nutritional condition drives plasticity in biting time of day, suggesting it is an underappreciated contributor to residual malaria transmission. Furthermore, our results suggest that targeting mosquito nutrition (e.g. sugar-baited traps) could influence mosquito behaviour in ways that affect the success of ITNs. More broadly, understanding the drivers of biting time of day variation is crucial for the future success of vector control tools and controlling malaria transmission.

## Background

The transmission of vector-borne diseases (VBDs), such as malaria and dengue, relies on the foraging rhythms of insect vectors, which take up pathogens from an infected vertebrate host during a blood meal and transmit them to a new host during a subsequent blood-feeding event (1,2). For example, *Plasmodium* parasites, the causative agent of malaria, are vectored between humans by *Anopheles* spp. mosquitoes, which are generally nocturnally active and preferentially bite between the hours of 11pm and 4am (*i.e.* the ‘classical’ time window) (3–5). Insecticide-treated bed nets (ITNs) exploit these rhythmic foraging behaviours, and are consequently one of the most effective methods for curbing the spread of malaria (6,7), averting 68% of deaths since 2000 (8). Despite this success, residual transmission occurs, in part due to the evolution of physiological insecticide resistance (e.g. biochemical and morphological modifications) and behavioural resistance in vector populations (9). In particular, millions of clinical malaria cases are predicted to be occurring annually due to mosquitoes altering the time of day they bite (10).

There are increasing reports that *Anopheles* spp. are shifting their biting time of day to earlier in the evening (‘early’ biting) or later in the morning (‘late’ biting) when humans are unprotected by ITNs (4,5,11–15), undermining the success of ITNs, especially in high-coverage regions. However, the extent to which mosquitos can shift biting time of day and the range of potential drivers are poorly understood. *Anopheles* spp. biting rhythms are weakly heritable, so shifts in biting time of day are expected to be predominantly driven by phenotypic plasticity (16). Phenotypic plasticity evolves to enable organisms to alter (via epigenetic mechanisms, for example) aspects of phenotype, including behaviours, in manners that maximise fitness in response to environmental variation (17).

Furthermore, variation in physiological condition and resource limitation can cause animals to alter their foraging rhythms (18) in adaptive (i.e. fitness enhancing) ways (19). For example, low food availability promotes a shift to diurnal foraging patterns in small nocturnal rodents, which minimises energy loss by remaining in burrows during particularly cold nights (20,21). Similarly, mosquitoes may garner greater benefits from shifting biting time of day when they are in poor nutritional condition, because the fitness benefits associated with acquiring a blood meal at suboptimal times of day outweigh the increased risks of desiccation and predation. Such context-dependent variation in the costs and benefits of plastic shifts in biting time may explain why biting times have seemingly not significantly changed in some *Anopheles* spp. populations despite high ITN use (22,23), and why the evolution of behavioural resistance varies across populations in manners that correlate with the abundance of sources of nutrition for mosquitoes (24).

Mosquitoes rely on blood and sugar meals for reproduction and survival, with female mosquitoes utilising proteins in blood for egg production, and using sugar sources to accumulate energy reserves for powering energetically demanding processes, such as flight (25). Their feeding ecology is under circadian clock control (26), with females seeking blood predominantly at night and approximately every 3 days to fuel gonotrophic cycles of egg development and oviposition (27), and feeding on sugar sources in between these cycles to promote survival until future biting opportunities (28,29).

Mosquitoes are likely to vary in their nutritional status because the type and abundance of sugar sources available to mosquitoes varies across habitats and seasons (25,30), and vector control tools such as ITNs can limit access to protein rich blood meals (31). In response to variation in food sources, mosquitoes exhibit plasticity in multiple feeding behaviours. If a preferred host is not found, mosquitoes are more likely to feed on sugar (32) or choose an alternative host (23), and will feed on low-sugar plant tissues when sugar-rich plants are unavailable (33). However, despite the importance of mosquito foraging rhythms for malaria transmission and that physiological condition alters foraging rhythms in other taxa, the effect of nutritional resource availability on mosquito biting time of day is unknown.

Here, we perturb the nutritional resources provided to adult mosquitoes to test whether nutritional constraints cause a shift in biting time of day. Using a semi-field system, we simulated scenarios in which mosquitoes have taken an initial blood meal (simulating the point at which malaria parasites would be acquired) and then resided in environments that provide varying access to further blood meals and sugar concentrations. Our three scenarios represented mosquitos that: (i) were unsuccessful at acquiring a subsequent blood meal and only have access to low levels of sugar (0.5% sucrose), (ii) were unsuccessful at acquiring a subsequent blood meal but have access to high levels of sugar (10% sucrose), and (iii) successfully acquired a subsequent blood meal and have access to high levels of sugar (10% sucrose plus an additional blood meal). We hypothesised that nutritionally deprived mosquitoes, provided with limited sugar and blood, would be more likely to host seek and shift their biting behaviour to earlier or later in the diel cycle, compared to better-resourced mosquitoes. We also supplemented our data with a preliminary field collection to investigate if biting time of day of wild-caught adult mosquitoes correlates with nutritional condition. We discuss the potential outcomes of shifts in biting time of day in response to resource availability for understanding residual transmission patterns, and how this may impact parasite development and transmission between hosts.

## Methods

We performed a semi-field experiment to assess if mosquito nutritional status impacts host seeking at different times of day. We reared wild-caught larvae, and maintained blood fed female *Anopheles gambiae s.l.* mosquitoes from the F_2_ generation on diets that varied in nutritional resources, before releasing them into an enclosed semi-field system with traps mimicking human odour. The three traps were programmed to separately capture mosquitoes biting at early (evening), classical (during the night) and late (morning) times of day. In addition, we collected wild adult *Anopheles* mosquitoes over the course of one night, to test for correlations between mosquito nutritional status and the time of day they attempted to bite.

### Study sites

For the semi-field experiment, we collected *Anopheles gambiae s.l.* larvae from three urban sites across the Greater Accra region of Ghana (Tuba, 5° 30’ 47’ N 0° 23’ 16’ W; Teshie, 5° 35′ 0″ N, 0° 6′ 0″ W; East Legon, 5° 38’ 16.39’ N, 0° 9’ 40.33’ W) during the dry season in Dec 2023-Jan 2024 (Additional file 1, Fig. S1). Adult *An. gambiae s.l.* collections were conducted in Teshie during the dry season in February 2024. The most abundant malaria vector species in these three sites are *An. gambiae s.s*. and *An. coluzzii*, which both have a high Human Biting Index and predominantly bite between 10pm and 4am (34,35). We transported larvae to the insectary at the Department of Medical Microbiology, University of Ghana Medical School, Accra, and raised them to adults. We conducted mosquito behavioural experiments in January-February 2024 in an enclosed semi-field system (the “MalariaSphere”, (38,39)) at the University of Ghana campus in Legon, Accra. The average (± SEM) minimum and maximum daily temperatures during the experimental releases were 25.8±0.6°C and 34.5±0.4°C respectively (40). The enclosure was 5.8 x 4.2 x 2.8m and covered in an insect-proof screen to prevent mosquito escape and/or entry from the external environment.

### Semi-field host seeking experiment

To ensure sufficient sample sizes, we conducted semi-field experiments using the F_2_ progeny of wild-caught *An. gambiae s.l.* larvae (Fig 1A). We pooled collected larvae and then transferred them into large tubs of water. We kept larvae at similar densities throughout the rearing procedure and fed them on TetraMin Baby fish food. The emerged adults formed the F_0_ generation, which we maintained in cages under standard rearing conditions (26±2°C, 80% relative humidity with a 12L:12D cycle), with *ad libitum* access to 10% sucrose solution in water. We provided a blood meal via direct-feeding (36), and placed soaked cotton wool in petri dishes into the cages for egg laying. We reared the resulting F_1_ generation larvae and adults under the same conditions, and their eggs were used to produce the experimental F_2_ generation. On day 1 post-emergence of F_2_ adults, we randomly allocated female mosquitoes to three experimental cages (n=100-150 per cage) and allocated each cage to one of three feeding treatments (Fig.1A): (i) 0.5% sucrose in water (0.5% suc), (ii) 10% sucrose in water (10% suc) or (iii) 10% sucrose in water plus an additional blood meal on day 6 post-emergence (10% suc + bm). We maintained all treatment groups on their respective sucrose solutions *ad libitum* from day 1 post-emergence until the host seeking experiment. In addition, to mimic mosquito life history in nature, we provided all groups with an initial blood meal on day 3 post-emergence. We gave mosquitoes the opportunity to lay eggs after all blood meals. On day 9-10 post-emergence, we transferred mosquitoes from each cage to paper cups (n=70-100 per cup), where they were housed for six hours with access to water only prior to assaying biting time of day. We reared six batches of mosquitoes to conduct a total of six semi-field system releases.

**Figure 1.**
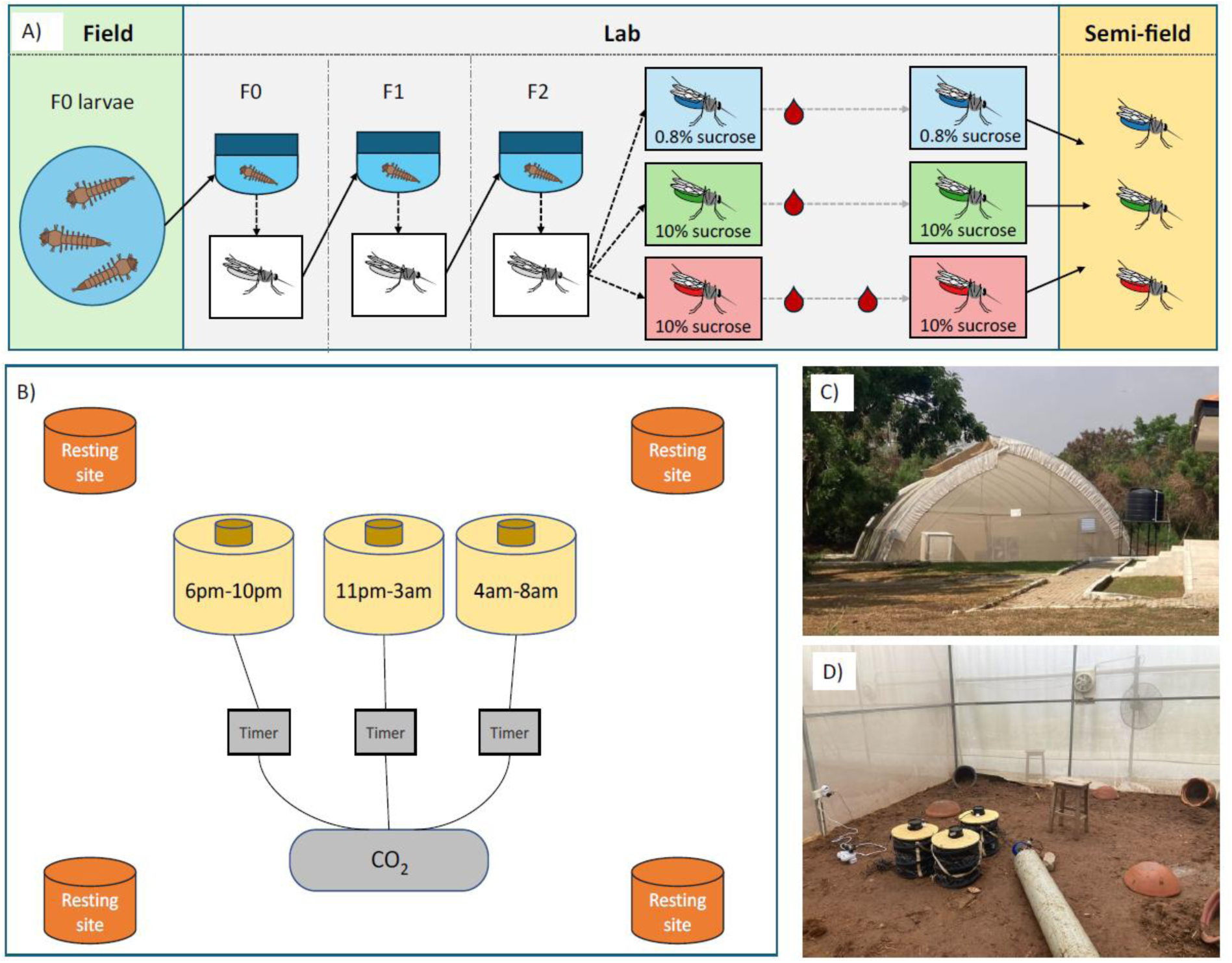
The experimental design for the semi-field behavioural experiment. Schematic diagrams of the entire experiment, from wild larvae collection to release of mosquitoes reared under the three feeding treatments into the semi-field enclosure (A), and the semi-field assay set-up (B), featuring four resting sites for mosquitoes, and traps baited with human odour which were programmed to turn on at 6pm-10pm, 11pm-3am and 4am-8pm respectively. These windows correspond to approximately ZT12-ZT16, ZT17-ZT21, and ZT22-ZT2 respectively, where Zeitgeber time (ZT) defines the hours since lights on and ZT0/24 is dawn. Each trap was linked to the CO_2_ source, using separate BG-CO_2_ timers to ensure the CO_2_ was only flowing when the corresponding trap that was switched on. Photographs show the semi-field facility (C) and the behavioural assay set-up (D).

We placed three BG-Sentinel traps (Biogents, Germany) in the semi-field enclosure, with an MB5 human-mimic lure developed to attract anopheline mosquitoes (Biogents, Germany) and a CO_2_ flow of approximately 21g/hour, to mimic human scent and breath. We programmed each trap to turn on at either 6pm-10pm to capture ‘early’ biters, 11pm-3am to capture ‘classical’ biters, or 4am-8am to capture ‘late’ biters (Fig. 1B-D). We chose these timings because mosquitoes in this region predominantly bite between 11pm and 4am, but biting can occur from approximately 6pm and persist into the morning beyond 6am (34). Thus, these time windows also correspond to when people are typically indoors and protected by ITNs (classical), or are outdoors and unprotected (early and late) (37). CO_2_ flow was controlled with BG-CO_2_ timers (Biogents, Germany), ensuring CO_2_ was only emitted when the respective trap was switched on.

Within the MalariaSphere, released mosquitoes could access resting sites (four clay pots around the enclosure). We did not provide water or food sources because this would have confounded the experimental treatment groups. We colour-marked mosquitoes from each feeding treatment using aerosolised fluorescent powder (FTX series, Swada London), applied following Machani et al (38). We rotated the colour of each treatment group across sequential releases to ensure any differential impacts of colour on mosquito behaviour did not confound treatment groups. We carried out six releases over a period of three weeks, with mosquitoes being released into the enclosure between 5pm-5:30pm, just prior to dusk at 6pm (n=235-300 mosquitoes per release). A period of at least 48 hours between each release and checking all resting sites with a Prokopack aspirator ensured that any remaining untrapped mosquitoes in the MalariaSphere had been removed or died before the next release. We collected mosquitoes from the traps, at 8am the day after the release, then cold-anaesthetised and counted the number caught in each trap.

### Wild mosquito collections

In addition to the experiment, we also conducted a preliminary investigation into correlations between biting time of day and nutritional content in wild mosquitoes, by collecting adult mosquitoes (n=95) overnight between 6pm and 7am using the human landing catch (HLC) technique. This technique involves trained human collectors exposing their lower legs and collecting mosquitoes that land on them before they bite, and is the gold standard for investigating human exposure to disease vectors (39). Every hour, we snap-froze collected mosquitoes on dry ice and identified specimens to genus level according to morphological characteristics (40), noting the number of *Anopheles* spp. mosquitoes collected per hourly bin. We placed individual anopheline mosquitoes in 1.5ml microcentrifuge tubes and stored them on dry ice, transferring them to a −20°C freezer after the overnight collection period had ended.

### Nutrition assays

Immediately prior to each semi-field release in the experiment, we collected a subset of mosquitoes from each feeding treatment group (n=14-19 per group, total n=48) to confirm our resource perturbations impacted nutritional status. We also quantified mosquito nutritional status in individual wild-caught mosquitoes (n=66). Mosquitoes were frozen at −20°C for lipid, glycogen and total sugar analyses using modified Van Handel protocols (41–43). Firstly, we removed a wing from each mosquito to quantify body size. Wing length is a commonly used proxy for adult body size because dry weight and wing length are highly correlated (44), and is included in statistical analyses to ensure any size variation does not confound differences in nutritional content. We lysed individual mosquitoes in 100µl 2% sodium sulphate and added 750µl of 1:2 chloroform:methanol. We centrifuged samples at 12000 rpm for 3 minutes; the supernatant was used for lipid and total sugars analysis, and the precipitate was used for glycogen analysis. We conducted lipid, glycogen and total sugars assays following Oke et al (45). For wild-caught mosquitoes, we used 250µl supernatant for total sugars analysis.

### Statistical analysis

We used R v. 4.1.3 to perform data analysis. We confirmed that adult body size did not differ across the treatments using a linear model with wing length, mosquito release batch and their interaction as main effects (Additional file 1, Fig. S2). To analyse nutritional content (lipid, glycogen, total sugars) of individual mosquitoes, we used linear mixed models with feeding treatment as a main effect and mosquito release batch as a random effect, accounting for any within treatment variation in body size. We used binomial generalised linear mixed models (glmm) with mosquito release as a random effect to account for any between-release variation to investigate how feeding treatment impacted the overall proportion of mosquitoes caught across the night, with feeding treatment as a main effect, and how feeding treatment impacted the proportion of mosquitoes caught across the different biting time of day windows, with feeding treatment, trap time and their interaction as main effects. We estimated proportion of caught mosquitoes and relative odds ratios (OR) ± SE from models using the *emmeans* (46) package. We square root transformed nutrition data to meet assumptions of normality and homogeneity of variance. We minimised all models using likelihood ratio tests and AICc for non-nested models, and confirmed model assumptions using the *easystats* (47) package. To account for differences between mosquito release batches, we present estimated marginal means ± SEM (*emmeans* package), predicted from models. Whenever the minimised model contained a significant effect of feeding treatment, we conducted post hoc pairwise comparisons using the Tukey method with the *emmeans* package.

To investigate how nutritional content between field-caught and lab-reared mosquitoes varied, we used linear models to compare between the field-caught, 0.5% suc, 10% suc and 10% suc + bm groups. We performed a chi-square test to investigate if the number of field-caught mosquitoes differed throughout the night by grouping hourly bins into ‘early’ biters (6pm-11pm), ‘classical’ biters (11pm-4am) and ‘late’ biters (4am-7am). Using linear models, we analysed the nutritional content (lipid, glycogen, total sugars) of individual wild-caught mosquitoes with biting time of day as an unordered factor, and assessed the correlation between wing length and nutritional content. Nutrition data were square root transformed to meet assumptions of normality and homogeneity of variance. We conducted post hoc pairwise comparisons using the Tukey method with the *emmeans* package to compare nutritional content between field-caught and lab-reared mosquitoes.

## Results

### Resource availability perturbs mosquito nutritional status

First, we confirmed that mosquitoes fed on the lowest concentration of sucrose (0.5%) had lower nutritional reserves than mosquitoes fed 10% sucrose with or without an additional blood meal. Mosquitoes fed on 0.5% sucrose had significantly lower lipid levels (55.5±17.8µg), approximately 2.5-fold lower than those fed with 10% sucrose (138±31.4µg) and 3.3-fold lower than those fed 10% sucrose plus an additional blood meal (182±29.6µg) (χ^2^_2_=17.6, p < 0.001; Fig. 2A, Table 1). A similar pattern was observed for total sugars: mosquitoes fed with 0.5% sucrose had the lowest total sugar levels (85.6±69.9µg), approximately 5-fold lower than those given 10% sucrose (420±158µg) and 10% sucrose plus an additional blood meal (439±156µg) (χ^2^_2_=30.8, p < 0.001; Fig. 2B, Table 1).

**Figure 2.**
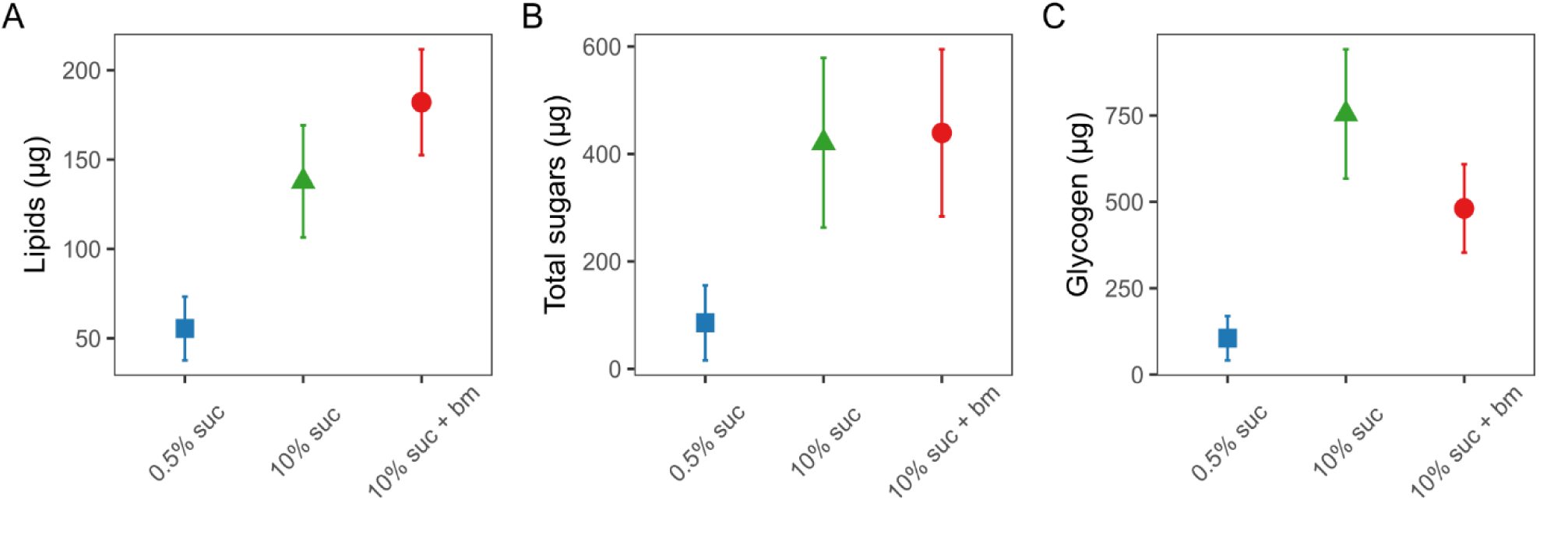
Concentrations (µg) per mosquito of lipids (A), total free sugars (B) and glycogen (C) in individual mosquitoes under differing feeding treatments on day 10 post-emergence. Nutritional perturbations began on day 1 post-emergence and all treatment groups received a blood meal on day 3 post-emergence (with an additional blood meal given to the 10% suc + bm group on day 6). Data presented are estimated marginal means ± SEM.

**Table 1.** Post hoc pairwise comparisons for nutritional content of mosquitoes provided with different feeding treatments. Significant p-values (<0.05) are highlighted in bold.

|  |  | *Test*<br>*statistic* | *p-value* |
| --- | --- | --- | --- |
| **Lipids** | 0.5% suc – 10% suc | $t = 2.75$ | **0.02** |
| | 0.5% suc – 10% suc + bm | $t = -4.54$ | **<0.001** |
| | 10% suc – 10% suc + bm | $t = -1.16$ | 0.48 |
| **Total sugars** | 0.5% suc – 10% suc | $t = 5.13$ | **<0.001** |
| | 0.5% suc – 10% suc + bm | $t = -6.13$ | **<0.001** |
| | 10% suc – 10% suc + bm | $t = -0.21$ | 0.98 |
| **Glycogen** | 0.5% suc – 10% suc | $t = 4.66$ | **<0.001** |
| | 0.5% suc – 10% suc + bm | $t = -3.69$ | **0.002** |
| | 10% suc – 10% suc + bm | $t = 1.55$ | 0.28 |

Finally, mosquitoes fed with 0.5% sucrose had the lowest glycogen levels (105±64.2µg), approximately 7-fold lower than those given 10% sucrose (754±187µg), with mosquitoes given 10% sucrose plus an additional blood meal exhibiting intermediate levels (481±128µg) (χ^2^_2_=21.7, p < 0.001; Fig. 2C, Table 1).

### Resource availability affects host seeking tendency and biting time of day

Across all six releases and trapping times of day, the total mosquitoes trapped ranged from 21% to 49%, with a mean of 32.4% of mosquitoes caught in the traps. The likelihood of host seeking at any time of day correlated negatively with the level of nutritional resources provided (χ^2^_2_=65.8, p < 0.001; Fig. 3A). Specifically, mosquitoes fed 0.5% sucrose were 2-fold more likely to be trapped than those fed 10% sucrose (odds ratio (OR): 2.08±0.28) and 3-fold more likely than those fed 10% sucrose plus an additional blood meal (OR: 2.97±0.42). Mosquitoes fed only 10% sucrose were 1.4-fold more likely to be trapped than those fed 10% sucrose plus an additional blood meal (OR: 1.43±0.21).

**Figure 3.**
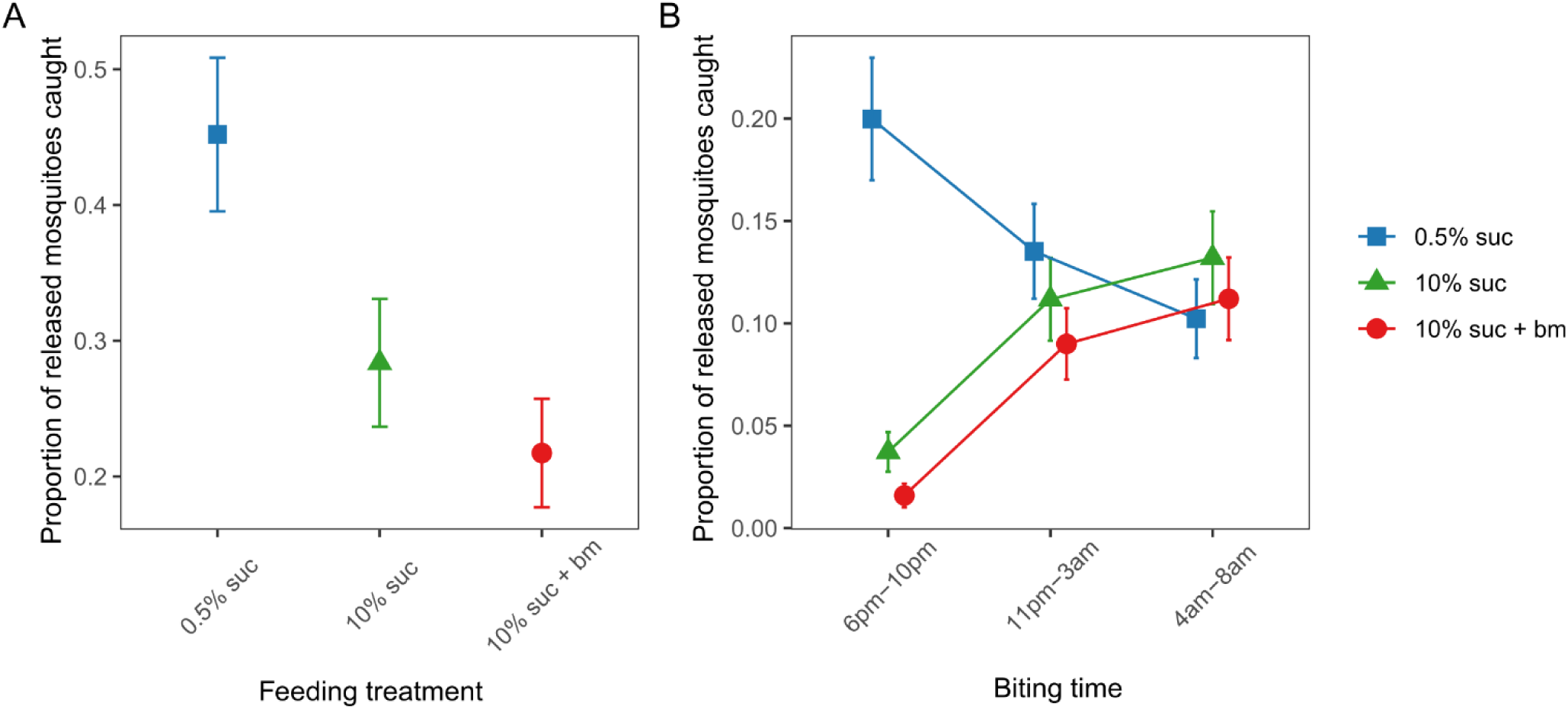
The proportion of released mosquitoes from each feeding treatment that were trapped across the entire night (A) and that were caught in the evening (early, 6pm-10pm), classical night time biting window (11pm-3am) and morning (late, 4am-8am) biting (B). Data presented are estimated marginal means ± SEM.

We also found that nutritional resources differentially affected the time of day of host seeking (interaction between treatment and trapping time: χ^2^_4_=95.4, p < 0.001; Fig 3B). The majority of early biting mosquitoes were from the 0.5% sucrose group, whereas mosquitoes with more resources were more likely to be caught at classic and late times of day. Specifically, mosquitoes fed 0.5% sucrose were approximately 5- and 10-fold more likely to be caught in the early biting window (20±2.9%) than those fed with 10% sucrose (3.7±0.9%) and 10% sucrose plus an additional blood meal (1.6±0.5%), respectively. At the classical and late trapping times of day, the proportion of mosquitoes fed 0.5% sucrose caught was similar to mosquitoes fed 10% sucrose with or without an additional blood meal. Mosquitoes fed 10% sucrose with or without an additional blood meal followed similar temporal patterns across all trapping times of day, with approximately 4-fold more of these mosquitoes being trapped during the classical (10% sucrose: 11±2.0%, 10% sucrose + bm: 9.0±1.7%) and late biting windows (10% sucrose: 13±2.3%, 10% sucrose + bm: 11±2.0%) than during the early biting window (10% sucrose: 3.7±0.9%, 10% sucrose + bm: 1.6±0.5%).

### Biting time distribution and nutritional status of wild host-seeking mosquitoes

We conducted a preliminary collection of wild host seeking *An. gambiae s.l.* mosquitoes to assess whether mosquito nutritional status correlates with biting time of day. We caught a total of 95 wild host seeking mosquitoes during our overnight collection, with a peak in biting between 12am-1am. We grouped hourly bins into ‘early’ biters (6pm-11pm), ‘classical’ biters (11pm-4am) and ‘late’ biters (4am-7am) and found that there were differences in the number of mosquitoes caught across these windows. Specifically, 68% of mosquitoes were caught during the classical biting window, compared to 7% and 24% caught during the evening and morning windows respectively, confirming that this population predominantly bite during the ‘classical’ time of day (χ^2^_2_= 45.0, p < 0.001; Fig 4).

**Figure 4.**
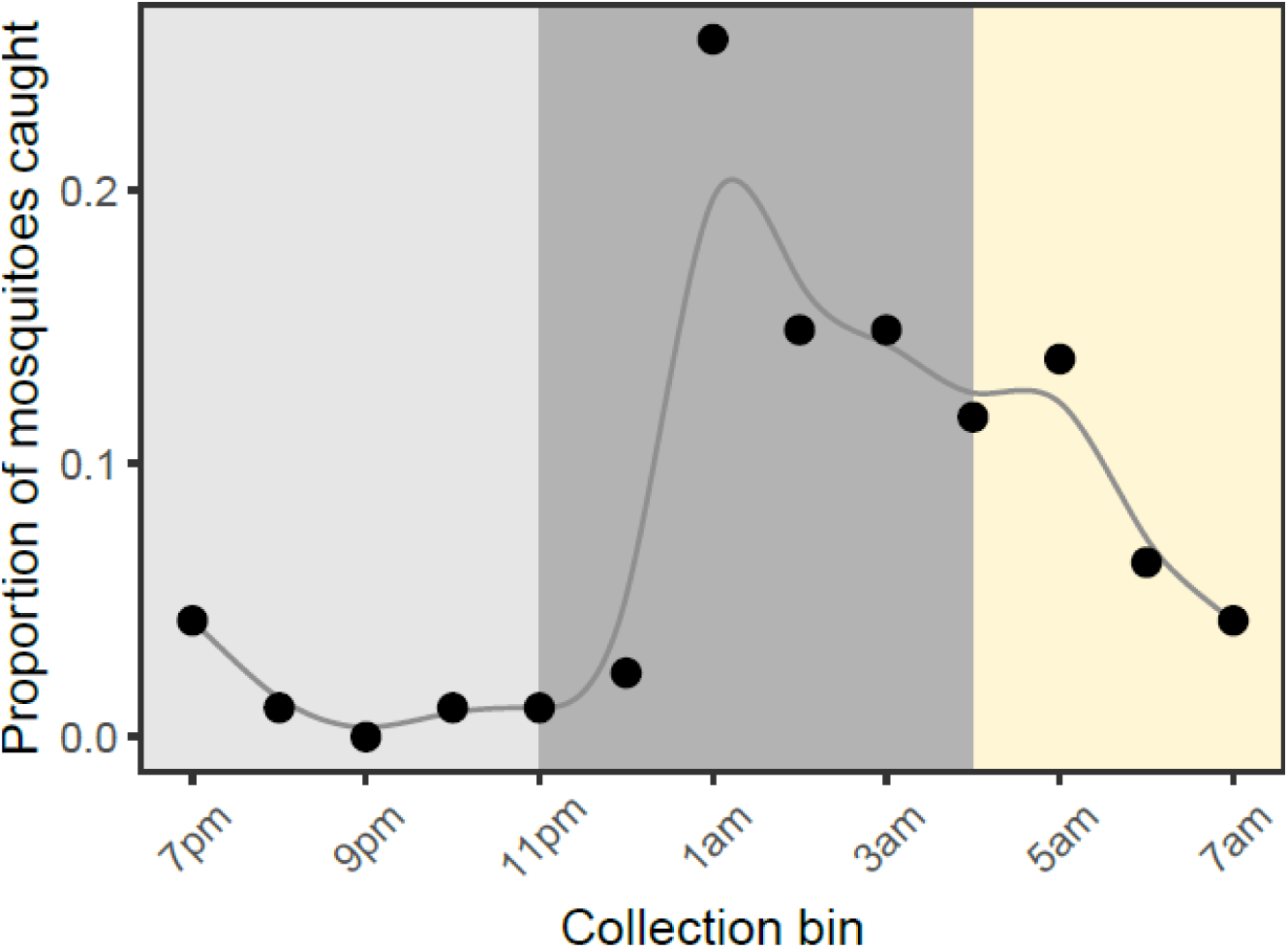
The proportion of wild adult *An. gambiae s.l.* mosquitoes caught by HLC collections across hourly bins between 6pm and 7am. Early, classical, and late bins are indicated by shaded regions from left to right. To highlight how the pattern changes over the course of the night, data points are connected by an X-spline.

Although wild mosquitoes varied in their nutritional content, we found no differences between lipid (F_9,56_ = 0.75, p = 0.66), sugars (F_9,56_ = 1.06, p = 0.40) or glycogen (F_9,56_ = 0.40, p= 0.93) contents across hourly bins (Additional file 1, Fig S3). However, we found that wild host-seeking mosquitoes had similar nutritional reserves to our experimental mosquitoes fed a 0.5% sucrose diet (estimated marginal means for field-caught mosquito lipids: 39.7±5.5μg, sugars: 65.2±13μg, and glycogen: 51.4±11μg), but had significantly lower reserves than mosquitoes fed 10% sucrose and 10% sucrose plus an additional blood meal (lipid: F_3,110_=24.8, p<0.001; sugars: F_3,110_=25.9, p<0.001; glycogen: F_3,110_=58.1, p<0.001; Table 2).

**Table 2.** Post hoc pairwise comparisons of the nutritional content of field-caught mosquitoes versus. lab-reared mosquitoes on different feeding treatments. Significant p-values (<0.05) are highlighted in bold.

|  |  | *Test statistic* | *p-value* |
| --- | --- | --- | --- |
| **Lipids** | field – 0.5% suc | $t = -1.26$ | 0.59 |
| | field – 10% suc | $t = -4.83$ | **$<0.001$** |
| | field – 10% suc + bm | $t = -7.93$ | **$<0.001$** |
| **Total sugars** | field – 0.5% suc | $t = -1.14$ | 0.67 |
| | field – 10% suc | $t = -4.96$ | **$<0.001$** |
| | field – 10% suc + bm | $t = -8.07$ | **$<0.001$** |
| **Glycogen** | field – 0.5% suc | $t = -1.97$ | 0.21 |
| | field – 10% suc | $t = -11.3$ | **$<0.001$** |
| | field – 10% suc + bm | $t = -8.87$ | **$<0.001$** |

## Discussion

By altering sugar provision and access to blood meals, and using traps that mimic human breath and scent, we find that resource availability modulates the time of day that *Anopheles gambiae s.l.* mosquitoes in Ghana search for a blood meal. Mosquitoes on a low sucrose diet had significantly lower levels of lipids, glycogen and sugar, and were 5-10 fold more likely to host seek in the evening (early) than mosquitoes on a high sucrose diet with or without a blood meal (Fig. 2, 3B). In contrast, better-resourced mosquitoes were most likely to host seek during the classical biting time window and into the morning (Fig. 2, 3B). We also find that as resources increased, the overall tendency to host seek decreased; mosquitoes on a low sucrose diet were 2-fold more likely to be caught compared to mosquitoes given a higher concentration of sucrose, and 3-fold more likely than those with access to 10% sucrose and an additional blood meal (Fig. 3A). Finally, to probe the ecological relevance of our experimental results, we also undertook a trapping session for wild host-seeking *An. gambiae s.l.* mosquitoes. While the nutritional content of wild mosquitoes varied, we found that levels were similar to the profiles of our low-resourced experimental mosquitoes fed 0.5% sucrose (Fig. 4). Given that mosquito flight during host seeking is energetically consuming (48), this suggests that our experimental feeding treatments resulted in mosquitoes with nutritional contents within an ecologically realistic range.

Foraging at unusual times of day can be detrimental because it exposes organisms to rhythmic environmental risks, such as the active phase of predators and suboptimal temperature/humidity, and misalignment between metabolic and other circadian rhythms may impair digestion and metabolism (1). However, under nutritional stress, finding food becomes more critical for survival, outweighing the environmental risks and physiological costs (18,21). For example, a poorly-resourced mosquito may begin host seeking earlier in the night because their lack of energy reserves reduce the chance of surviving until later at night (or the morning, depending on whether ITNs limit access to hosts at night). Our results are consistent with this hypothesis because poorly-resourced mosquitoes with low nutritional reserves are more likely to host seek, predominantly in the early evening when temperatures are higher and humidity is lower, and they risk being killed because human hosts are alert. Although we did not quantify survival, we did not encounter any live mosquitoes fed on a 0.5% sucrose diet whilst clearing the semi-field systems the day after each release, but regularly encountered live mosquitoes fed on higher sucrose solutions and/or an additional blood meal. This indicates that mosquitoes fed 0.5% sucrose were low-resourced and did not have sufficient reserves to survive until morning. Overall, our results suggest that early biting could be an adaptive plastic temporal shift to maximise fitness in sugar-poor regions and where hosts are not easily accessible due to high ITN use.

Better-resourced mosquitoes on high sucrose diets (with or without an additional blood meal) had higher nutritional reserves and waited until the classical time window to host seek, in alignment with their circadian physiology and when environmental risks are likely to be lower, which also supports our hypothesis. However, we also found that better-resourced mosquitoes were just as likely as poorly resourced mosquitoes to be trapped later in the morning. Such late biting by well-fed mosquitoes was unexpected but may be due to the lack of water/food sources provided in the semi-field enclosures, forcing untrapped mosquitoes to host seek to mitigate dehydration caused by the rise in temperature and the reduction in humidity in the morning (49,50). Alternatively, well-resourced mosquitoes which had failed to locate a host (*i.e.* had not entered a trap) during the night may have become resource-depleted because flight is energetically demanding (48), leading to an increased propensity to continue host seeking because they do not have sufficient energy to survive until the subsequent night. Regardless of the explanation(s), our results reveal that adult nutritional condition drives variation in biting time of day.

Our semi-field system allowed mosquitoes to experience ecologically realistic temperature and humidity fluctuations and provided sufficient flight space, but excluded other ecological factors, including predation and access to alternative food sources. Thus, to fully investigate how resource availability impacts biting time, future studies should consider adding plants of varying sugar contents and alternative blood hosts into semi-field systems. However, these omissions were necessary in our experiment to isolate the effect of adult nutritional status on biting behaviour. To probe the ecological relevance of our experimental results, we also undertook a preliminarily trapping session for wild host-seeking *Anopheles gambiae* s.l. mosquitoes. We found that they had similar nutritional profiles to our low-resourced mosquitoes fed on 0.5% sucrose, suggesting that our experimental perturbations represented natural conditions. The low sample size for wild caught mosquitoes precludes comparing nutritional reserves between host seeking times of day but does suggest that adult nutritional content does not correlate with body size (a product of larval condition) (Additional file 1, Fig. S4). Thus, catching lowly resourced mosquitoes at the time of host seeking is consistent with our experimental results revealing that adult nutrition influences biting rhythms. Further field studies are necessary to build on our findings, including sampling host-seeking mosquitoes across multiple sites, both indoors and outdoors, throughout a 24-hour period, examining their nutritional status, and investigating when mosquitoes become active.

Like previous studies across Africa (3,35,51), we found a large proportion of wild mosquitoes (approximately 20%) host seeking during the morning when humans are unprotected by ITNs. This pattern of ‘late’ biting has become more pronounced since the widespread use of ITNs. Because biting time shifts have a genetic component (16), and universal coverage of ITNs is approximately 75% in Ghana (52), late biting could be an evasion strategy evolving in the mosquito population where the larval collections for our experiment were carried out, rather than an entirely plastic behavioural change. Further work is needed to ascertain the extent to which biting time of day is genetically determined and/or due to behavioural plasticity, because plasticity itself evolves and can facilitate or constrain evolution that genetically segregates mosquito populations into early, classical or late biters. Modelling suggests that early biting is more likely to evolve in sugar-poor environments than sugar-rich environments, for example, due to differences in the abundance and diversity of plants (24). However, in areas of fluctuating resource availability or ITN coverage, plastic genotypes that can switch between early, classical, or late biting depending on their environmental conditions should have higher fitness (17).

Both our semi-field experiment and preliminary field data reveal significant variation in biting time of day, and intuition suggests that early biting by poorly-resourced mosquitoes could sustain malaria transmission. However, the overall impact of shifts in mosquito biting time of day for malaria transmission is difficult to predict, because of complex and potentially opposing effects on both mosquito and parasite fitness. While early biting causes greater contact between vectors and hosts in regions with high ITN coverage, malaria parasite development is less productive in low-resourced mosquitoes, and mosquito survival is reduced (45). Thus, the poor nutritional state of early biters could curtail onward transmission through lower parasite numbers (53) and because vectors may not survive long enough for malaria parasites to complete their development (54,55). In addition, parasite establishment in mosquitoes is impacted by daily temperature rhythms (56) and parasite infectivity and mosquito susceptibility vary between day and night (1,57,58). Disruption to foraging rhythms also reduces fitness in other insects (59), and in mosquitoes could lead to mismatch between metabolic processes and blood digestion (60). The net outcome for malaria transmission of these myriad and potentially synergistic and antagonistic effects of biting time of day is challenging to predict. However, recognising that transmission is time of day dependent is necessary for accurately predicting epidemiology, the risk of clinical malaria cases, and the trajectory of parasite evolution.

### Conclusions

Our results suggest that plasticity in biting time of day, driven by variation in adult mosquito nutritional condition, may be an underappreciated contributor to residual malaria transmission. In particular, nutritional stress in mosquito populations may lead to shifts in biting behaviour, reducing ITN efficacy. This highlights that mosquito nutrition and its influence on the timing of biting behaviour should be considered when implementing and assessing vector control strategies, especially those which target mosquito foraging behaviour, such as ITNs and sugar-baited traps. More broadly, investigating the drivers of variation in biting time of day and the overall impact on pathogen transmission is critical for assessing and improving the efficacy of current and future vector control tools. This is especially important for malaria, where up to 30% of biting is reported to occur when humans are unprotected by ITNs and is predicted to lead to millions of additional clinical cases per year (3,10). However, we also expect our findings to apply to other vector-borne diseases that rely on vector foraging rhythms for between-host transmission.

## Declarations

### Ethics approval and consent to participate

The study protocols for mosquito colony blood feeds and human landing catches (HLCs) were ethically reviewed and approved by the Ghana Health Service Ethics Review Committee (GHS-ERC: 021/07/23). Volunteers involved in colony blood feeds were tested for malaria prior to direct-feeding, and the same person fed the mosquitoes throughout the experiment. Permission to conduct HLCs in study sites was obtained from the District Assembly Representative, and written consent was obtained from all collectors. Collectors had access to malaria prophylaxis and did not report any adverse events.

### Consent for publication

Not applicable.

### Availability of data and materials

The datasets supporting the conclusions of this article are available in the Edinburgh DataShare repository and can be accessed with [https://doi.org/LINK AVAILABLE UPON ACCEPTANCE].

### Competing interests

The authors declare that they have no competing interests.

### Funding

This work was funded by a Varley-Gradwell Travelling Fellowship in Insect Ecology (2023–2024) to C.E.O, and by a Wellcome Trust Investigator Award (202769/Z/16/Z) and Wellcome Trust Discovery Award (227904/Z/23/Z) to S.E.R. Research equipment and travel funds for M.G.M. and S.S.C.R. were funded by an Initiation Grant from the University of Notre Dame, Notre Dame Research office. Y.A. A. is funded by the National Institute for Health (D43 TW 011513). The funders had no role or influence on the design of this study, data collection, analyses, and interpretation of the data collected, as well as in writing this manuscript.

### Authors’ contributions

Conceptualisation: CEO, SSCR, YAA, SER. Methodology: CEO, SSCR, MGM, YAA, SER. Investigation: CEO, SSCR, MGM, ARMS, YAB. Analysis: CEO. Writing-original draft: CEO, SER. Writing-review and editing: CEO, SSCR, MGM, YAA, SER. All authors read and approved the final manuscript.

## Acknowledgements

We thank Anisa Abdulai for experimental and technical assistance, and the serology lab team at the Noguchi Memorial Institute for Medical Research for kindly letting us use their facilities.

## Additional files

Additional file 1 (.pdf) Supplementary Figures.

Figure S1 is a map of larval and adult collection sites, Figure S2 is the wing length of lab-reared mosquitoes under different feeding treatments, Figure S3 provides information about the nutritional contents of wild adult mosquitoes collected at different times of day, and Figure S4 is the correlation between wing length and nutritional content in wild adult mosquitoes.

## Notes

### Competing Interest Statement

The authors have declared no competing interest.

### Summary of Updates

Manuscript has been updated after first round of peer-review.

