## Additional file 1 for "Biting time of day in malaria mosquitoes is modulated by nutritional status"

### Supplementary Figures

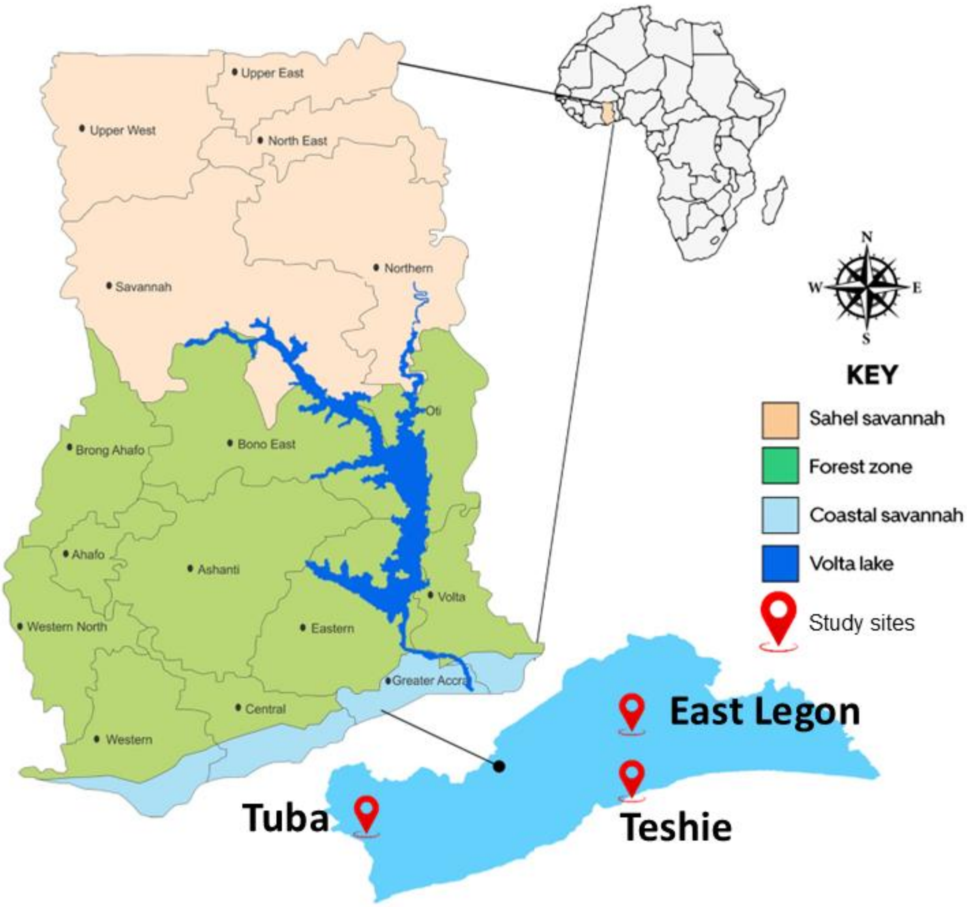


**Figure S1.** Map of larval and adult mosquito collection sites in Greater Accra, Ghana. Larvae from the three sites were collected in Dec 2023-Jan 2024, combined, and female $F_2$ progeny were used for mosquito nutrition perturbations and biting time of day assays. Wild adult mosquitoes were collected from Teshie across one night.

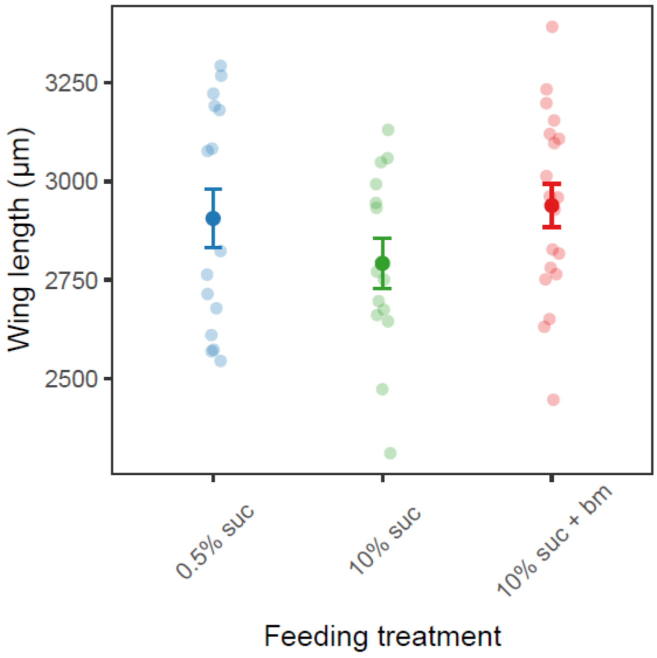


**Figure S2.** Wing length (a proxy for body size) of lab-reared mosquitoes under different feeding treatments. Data presented are means ± SEMs, with each point representing an individual mosquito. While mosquito size differed between release batches ($F_{2,45}$=5.55, p=0.007), this was consistent across feeding treatments within each batch ($F_{3,40}$=0.05, p=0.98). Overall, there was no difference between wing length across feeding treatment groups ($F_{2,43}$=0.66, p=0.52).

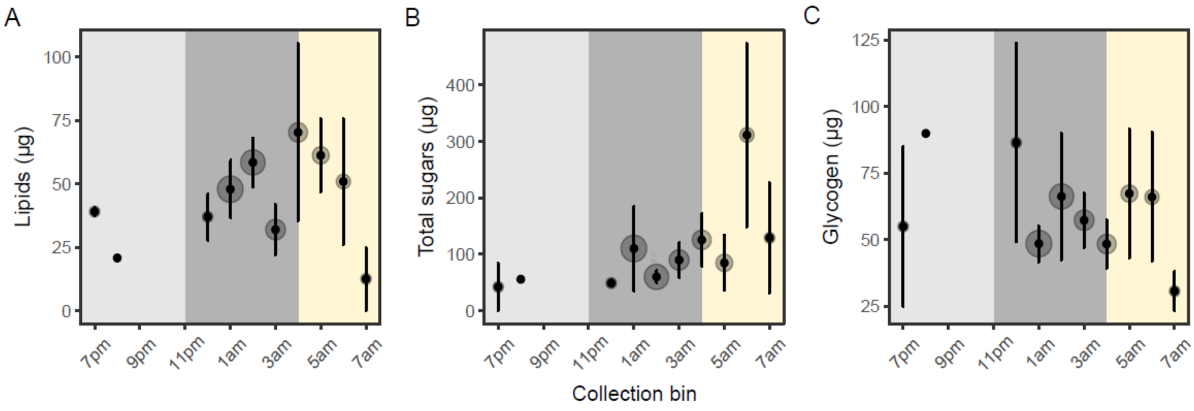


**Figure S3.** Concentrations (µg) per mosquito of lipids (A), total free sugars (B) and glycogen (C). Data presented are means ± SEM, and the size of the grey circle reflects the sample size per hourly bin.

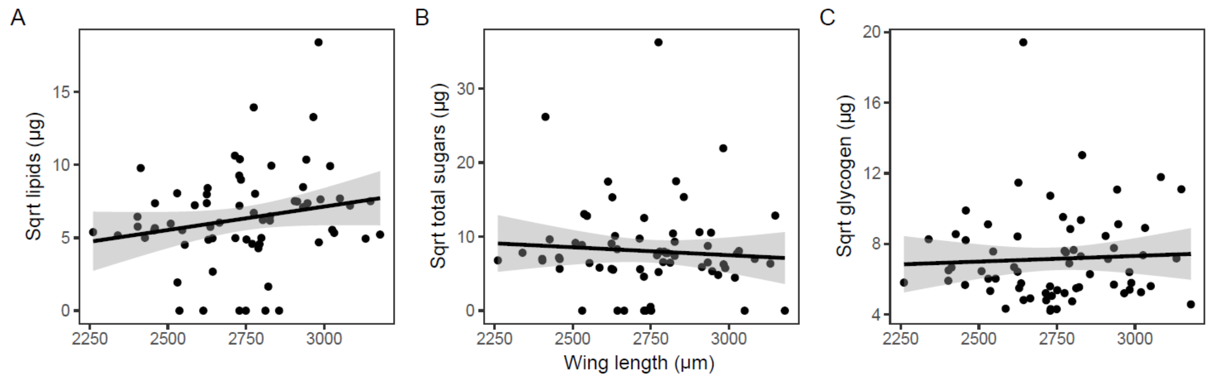


**Figure S4.** The relationship between wing length (a proxy for body size) and (A) lipids, (B) total free sugars and (C) glycogen in wild-caught adult mosquitoes. Data points represent an individual mosquito, with lines and shading denoting the best fit linear model estimates and 95% confidence intervals. For lipids ($F_{(1,64)}$ = 2.80, p = 0.10), total free sugars ($F_{(1,64)}$ = 0.35, p = 0.56) and glycogen ($F_{(1,64)}$ = 0.18, p = 0.67), there was no correlation with wing length.
